# Non-targeted metabolomics identifies erythronate accumulation in cancer cells

**DOI:** 10.1101/2022.12.04.519010

**Authors:** Jie Zhang, Mark A. Keibler, Wentao Dong, Jenny Ghelfi, Thekla Cordes, Tamara Kanashova, Arnaud Pailot, Carole Linster, Gunnar Dittmar, Christian M. Metallo, Tim Lautenschlaeger, Karsten Hiller, Gregory Stephanopoulos

## Abstract

Using a non-targeted isotope-assisted metabolomics approach, we identified erythronate as a metabolite that accumulates in several human cancer cell lines. Erythronate has been reported to be a detoxification product derived from off-target glycolytic metabolism. We provide data supporting a possible alternative route to erythronate production involving the dephosphorylation of the pentose phosphate pathway intermediate erythrose-4-phosphate to form erythrose, followed by the oxidation of erythrose by an aldehyde dehydrogenase. Finally, we detected increased erythronate concentrations in tumors relative to adjacent normal tissues from lung cancer patients. These findings suggest the accumulation of erythronate to be an example of metabolic reprogramming in cancer cells, raising the possibility that elevated level of erythronate may serve as a biomarker of certain types of cancer.

## Introduction

Metabolic reprogramming to sustain elevated energy, reducing equivalent, and biosynthetic precursor production is now recognized as a hallmark of cancer cells ^1–6^. One consequence of this rewiring of cellular metabolism is the upregulated biosynthesis and accumulation of metabolites that are minimally synthesized under normal conditions ^7–10^. Identification of such metabolites can yield effective tumor biomarkers and also elucidate the altered biosynthetic network induced in cancer cells, both of which can lead to advances in diagnostics and therapeutics. Investigations of these metabolic networks with stable-isotope metabolomics typically rely on the availability of targets that can be followed through the use of labeling substrates ^11,12^. However, these targets depend on preconceived notions about metabolism to define a list of target metabolites. On the other hand, non-targeted metabolomics methods, which do not require *a priori* knowledge of target metabolites in a sample, are useful for identifying pathways that are selectively activated in cancer cells. As such, Non-targeted Tracer Fate Detection (NTFD), a stable-isotope-based method developed in our lab, is capable of addressing this problem by detecting all metabolites that are derived from a labeled metabolic precursor, regardless of previous identification ^13,14^.

Erythronate has been reported to derive from the pentose phosphate pathway (PPP) intermediate erythrose-4-phosphate (E4P). The most comprehensively described biosynthetic pathway of erythronate synthesis comprises oxidation of E4P to 4-phosphoerythronate (4PE) via off-target glyceraldehyde-3-phosphate dehydrogenase (GAPDH) activity ^15^ followed by dephosphorylation of 4PE by the widely conserved phosphoglycolate phosphatase (PGP) ^16^. In this pathway, erythronate is produced as a detoxification product, because 4PE can strongly inhibit 6-phosphogluconate dehydrogenase, an enzyme in the oxidative PPP. Although this pathway provides an elegant biochemical rationale for erythronate production, it remains possible that other biosynthetic routes exist to at least some appreciable degree.

Additionally, most work describing erythronate production in mammalian cells has been performed with erythrocytes ^16,17^, and to our knowledge, erythronate has not yet been explicitly identified in cancer cells. It has been reported that elevated concentrations of erythronate were associated with low transaldolase activity seen in transaldolase deficient patients ^18^, suggesting a connection between erythronate and transaldolase, a key enzyme in the non-oxidative branch of the PPP (non-oxPPP). Given the importance of transaldolase and non-oxPPP in cell proliferation and growth ^19^, erythronate may be of high relevance in diseases like transaldolase deficiency and cancer.

In this work, we applied NTFD to cancer cells and identified erythronate as a metabolite significantly labeled by [U-^13^C^6^]glucose. We characterized the production of erythronate and intermediate metabolites using cell cultures and isolated enzymes, and we used these measurements to propose a possible alternative biosynthetic pathway. We also observed the differential elevation of erythronate in both lung tumors and cancer cell lines, suggesting the possibility that erythronate can be used as a biomarker for cancer.

## Results

### Non-targeted [^13^C]glucose labeling analysis identifies erythronate in cancer cells

Application of NTFD to A549 lung carcinoma cells determined several metabolites that were labeled by [U-^13^C^6^]glucose. However, most of these metabolites could not be identified due to the lack of a comprehensive library. Among those present in our library yet not part of central carbon metabolism, one of the most strikingly labeled was erythronate (**Fig. 1A**). Despite the fact that its presence in human bodily fluids has been recognized for decades, the biological role of erythronate remains ambiguous and its biosynthetic pathway is uncharacterized in public metabolic pathway databases, such as KEGG, MetaCyc, and BioCyc. Gas chromatography coupled to mass spectrometry (GC-MS) showed that fragment ion 409, which contains all four carbon atoms of the trimethylsilyl (TMS)-erythronate derivative, was mainly abundant as the M+4 (413, four times labeled) mass isotopomer in samples of the above cell lines when cultured with [U-^13^C_6_]glucose (**Fig. 1A, Figure S1**), meaning that erythronate is synthesized from glucose. Erythronate extracted from cells cultured in [1,2-^13^C_2_], [1,3-^13^C_2_], [4-^13^C], and [6-^13^C]glucose displayed labeling patterns that indicated that it was derived primarily from C3-6 of glucose, consistent with previous reports that identified the PPP intermediate E4P as a precursor (**Fig. 1B, Fig. S2**).

**Figure 1.**
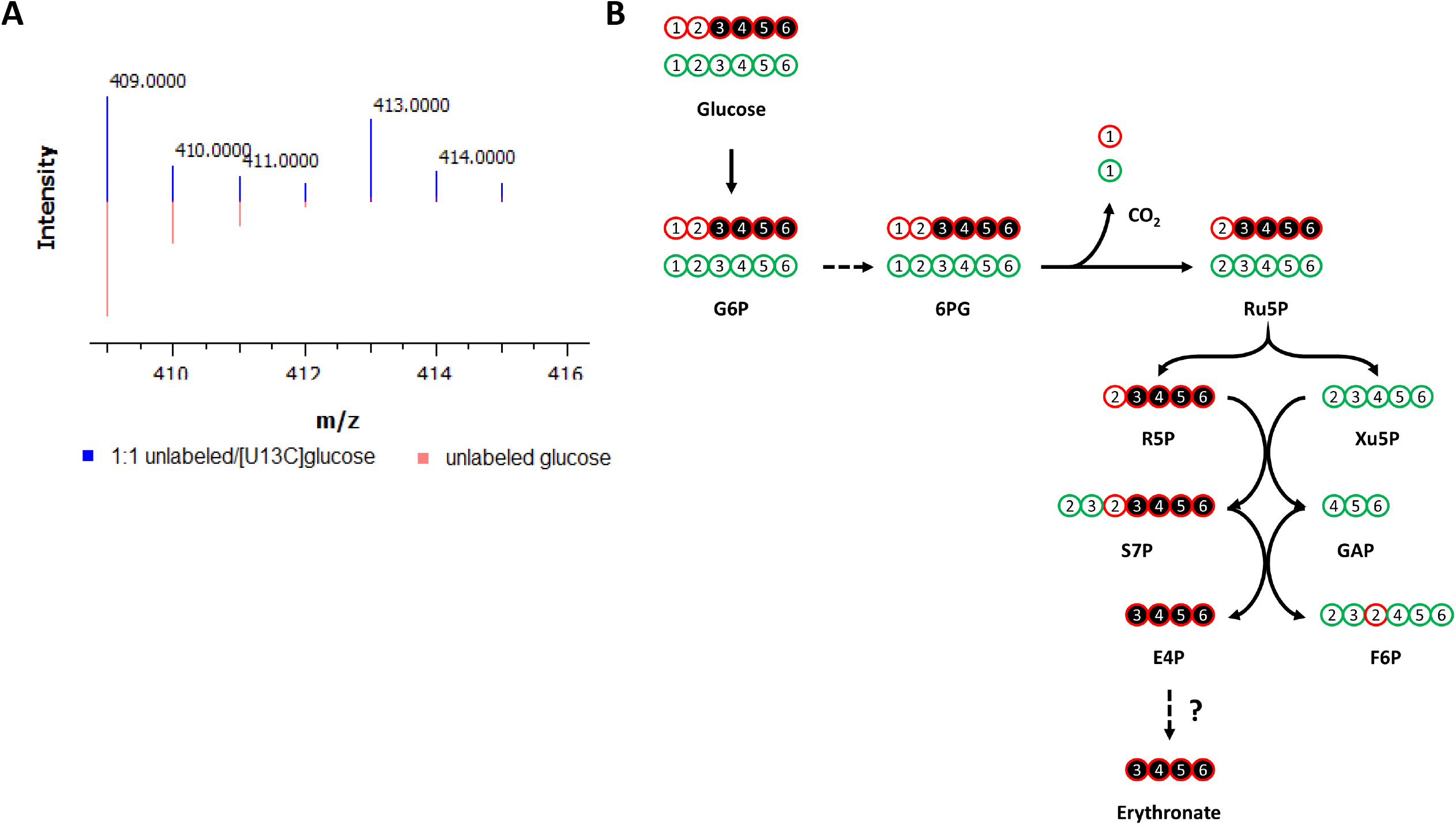
Labeling of metabolites by ^13^C-glucose in cancer cells. **A**) Labeling of erythronate by [U-^13^C_6_]glucose in cancer cells. A549 lung carcinoma cells were cultured with media containing either unlabeled glucose (red) or a 1:1 mixture of unlabeled and [U-^13^C_6_]glucose (blue). Intracellular polar metabolites were extracted from the cells and analyzed on GC-MS. The ion 409, which is one of the characteristic fragments of TMS-erythronate and contains all four carbon atoms of the molecule, is found to be M+4 labeled (m/z = 413), indicating that all four carbon atoms of erythronate are derived from glucose. **B**) Carbon transition map for [1,2-^13^C_2_]glucose through the pentose phosphate pathway. This map illustrates the fate of ^13^C-atoms (indicated bycircles) derived from [1,2-^13^C_2_]glucose, which is used to trace the metabolism of glucose *via* the pentose phosphate pathway (PPP) and glycolysis. The first ^13^C-atom of [1,2-^13^C_2_]glucose is lost through decarboxylation if G6P enters the oxidative branch of the PPP, but is retained if G6P is metabolized via glycolysis. Detailed chemical structures are shown in **Fig. S2**. G6P, glucose-6-phosphate; 6PG, 6-phosphogluconate; Ru5P, ribulose-5-phosphate; R5P, ribose-5-phosphate; Xu5P, xylylose-5-phosphate; F6P, fructose-6-phosphate; S7P, sedoheptulose-7-phosphate; GAP, glyceraldehyde-3-phosphate; E4P, erythrose-4-phosphate; CO_2_, carbon dioxide. This figure is adapted from Dong et al., 2019 ^56^.

### Enzymes involved in erythronate biosynthesis

To synthesize erythronate from E4P, two biochemical steps are required: dephosphorylation and oxidation of the aldehyde into its corresponding carboxylic acid. GAPDH has been documented to oxidize E4P to 4PE via off-target metabolism, and a more recent paper identifies PGP as the enzyme catalyzing dephosphorylation of E4P to erythronate ^16^. However, we speculated that other enzymes could also be partially responsible for erythronate production due to enzyme promiscuity ^20–22^, possibly with dephosphorylation occurring before oxidation.

We first looked for enzymes that can potentially catalyze the oxidation of the aldehyde group on erythrose. A549 cells cultured in DMEM supplemented with 1 g L^-1^ [U-^13^C_4_]erythrose produced high amounts of erythronate mainly as the M+4 isotopomer (**Fig. S3**), suggesting that erythrose can cross the cell membrane and be oxidized into erythronate. To assess cofactor dependency, A549 cell lysates were used in an *in vitro* erythrose oxidation assay in the presence of the common oxidizing cofactor nicotinamide adenine dinucleotide (NAD^+^). Analysis of light absorbance at 340 nm, the λ^max^ of NADH, and the reaction products by GC-MS showed that cell lysates could oxidize erythrose to erythronate in the presence of NAD^+^. This is consistent with the NAD^+^ requirement of E4P-to-4PE oxidation by GAPDH (**Fig. 2**).

**Figure 2.**
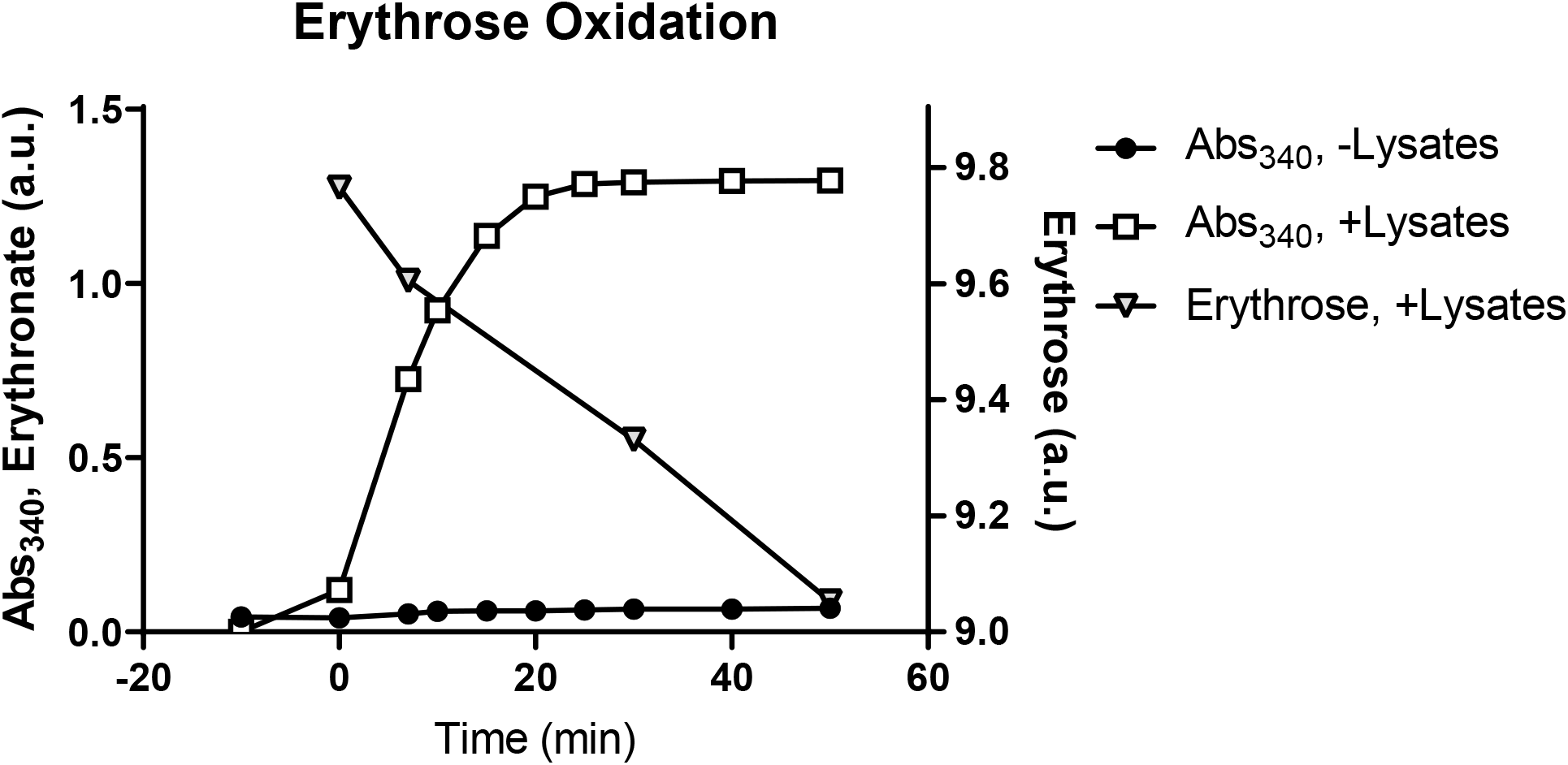
Conversion of erythrose to erythronate by A549 cell lysate. A549 cell lysates were used in an enzymatic assay that converts erythrose to erythronate in the presence of NAD^+^. The conversion was indicated by the formation of NADH, which can be monitored by absorption at 340 nm (Abs_340_). Samples were taken from the assay at various time points to determine the depletion of erythrose and the formation of erythronate.

To identify the enzyme(s) responsible for the oxidation of erythrose to erythronate, we performed protein fractionation coupled with the NADH light absorbance assay. The protein fraction conferring the highest erythrose-oxidizing activity was collected, and the identities of proteins present in this fraction were determined using Orbitrap liquid chromatography tandem mass spectrometry (LC-MS-MS). The most prominent hit was the aldehyde dehydrogenase 1 family member A1 (encoded by the gene *ALDH1A1* in humans). An *in vitro* enzyme assay with ALDH1A1 protein purified using a polyhistidine tag confirmed that this protein could readily convert erythrose to erythronate in the presence of NAD^+^ (**Fig. S4**). We also found that glyceraldehyde-3-phosphate dehydrogenase (GAPDH) showed activity in converting erythrose into erythronate (**Fig. S5**). After transfecting A549 cells with siRNA targeting *ALDH1A1*, we observed that a 10-fold reduction in the mRNA expression level lowered the intracellular erythronate concentration by about 60% (**Fig. 3A-B**), suggesting that this isoform of aldehyde dehydrogenase may be responsible for the majority of erythronate formation in this cell line.

**Figure 3.**
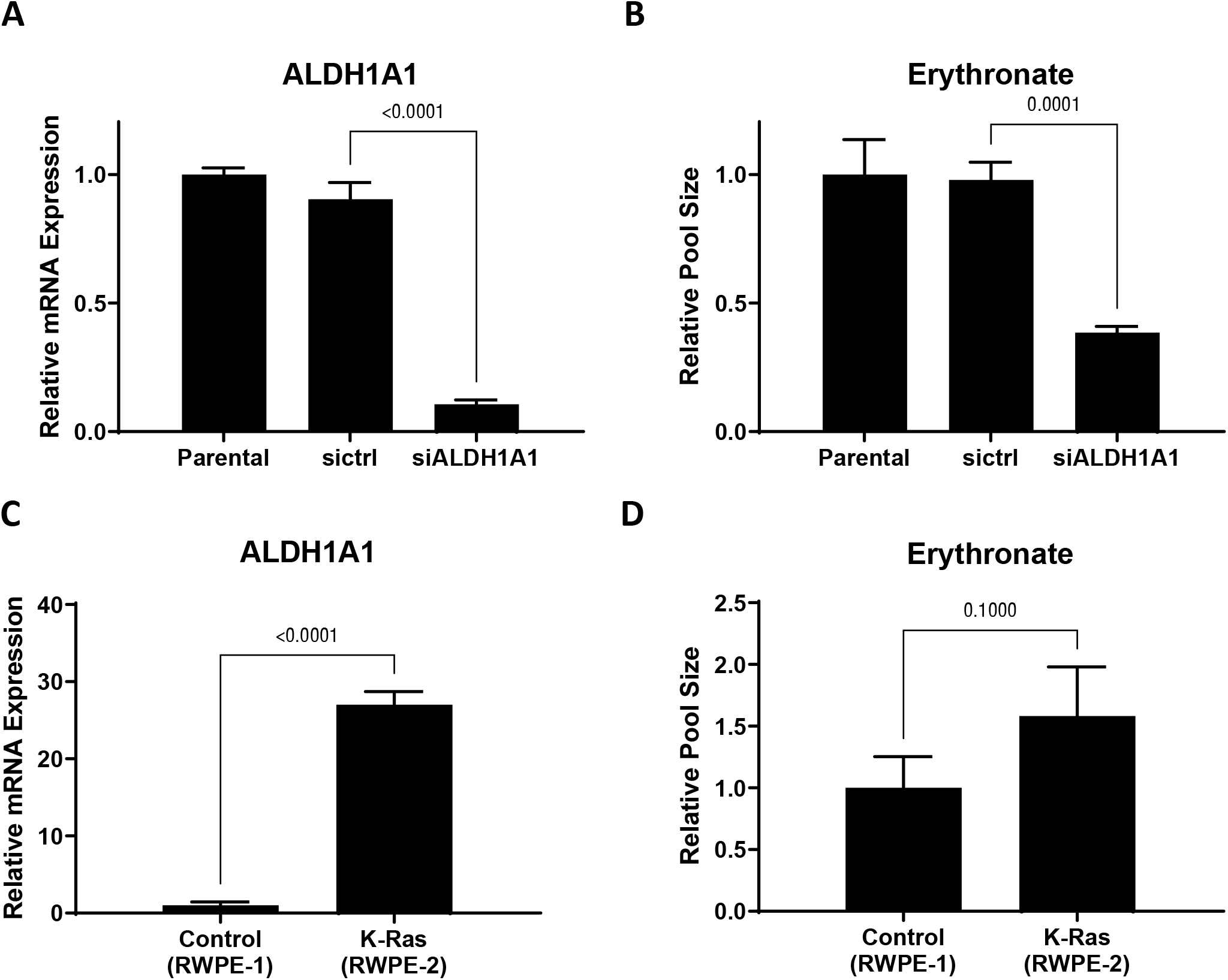
Changing *ALDH1A1* expression alters intracellular erythronate level. **A**) *ALDH1A1* mRNA was knocked-down by siRNA in A549 cells. **B**) *ALDHA1* knock-down led to substantially reduced erythronate levels. **C**-**D**) *ALDH1A1* expression level and the intracellular erythronate level in two human prostate epithelial cell lines. The two prostate epithelial cell lines RWPE-1 (non-cancerous) and RWPE-2 (tumorigenic) have very distinct *ALDH1A1* expression levels. Using the ribosomal protein L27 as the house-keeping gene, the mRNA level of *ALDH1A1* in RWPE-2 cells was determined to be 26-fold higher than that in RWPE-1 cells. Also, RWPE-2 cells produced approximately 60% more erythronate as compared to RWPE-1 cells. Error bars, s.d. (n=3). Statistical test: two-tailed paired t-test.

High aldehyde dehydrogenase activity has been considered as a marker of cancer stem cells ^23,24^, and ALDH1A1 has been suggested as a target for immunological therapeutics of many types of cancer ^25^. To further investigate the correlation between *ALDH1A1* expression and erythronate level, we used two prostate epithelial cell lines, RWPE-1, which is non-tumorigenic, and RWPE-2, a KRAS-transformed version of RWPE-1 that is capable of forming tumors in nude mice ^26^. *ALDH1A1* expression level in RWPE-2 cells was found to be 26-fold higher than that in RWPE-1 cells, and we observed 60% higher levels of intracellular erythronate in RWPE-2 cells compared to RWPE-1 cells (**Fig. 3C-D**). The correlation between *ALDH1A1* overexpression and higher erythronate level in the tumorigenic RWPE-2 cell line compared to the non-cancerous RWPE-1 cell line also suggests that high ALDH1A1 activity or erythronate concentration may have a role in cell tumorigenicity. This is consistent with several previous studies that suggest increased activity of ALDH1A1 is associated with tumorigenicity ^27,28^.

The other step required in the conversion of E4P to erythronate is the dephosphorylation of E4P to erythrose. The human genome encodes more than 200 phosphatases, most of which are classified as protein phosphatases ^29^. The most detailed biosynthetic pathway published for erythronate describes dephosphorylation of GAPDH-produced 4PE by the phosphatase PGP. We chose to test an acid phosphatase (AP), a type of abundant phosphatase identified in animals, plants and fungi, which has also been found to be relevant in the diagnosis and treatment of prostate cancer ^22^. Our results show that E4P was dephosphorylated and converted into erythrose under an acidic condition (pH∼4.8, **Fig. S6A**). This is consistent with the prior knowledge that the AP family is a type of highly promiscuous phosphatase ^22^ and demonstrates that AP is capable of generating erythrose from E4P.

Having demonstrated the NAD^+^-dependent oxidation and dephosphorylation in two separate reactions, we asked if E4P could be converted to erythronate. Since E4P cannot cross the cell membrane due to the negative charged phosphate group, we performed *in vitro* enzymatic assays using E4P as the substrate. Results showed clearly that A549 cell lysate could readily convert E4P into erythronate, and the overall reaction was significantly enhanced by the addition of 0.8 mM NAD^+^, confirming the NAD^+^ dependence of the oxidizing enzyme(s) (**Fig. S6B**).

### Effects of perturbation in the PPP on the erythronate level

We have demonstrated that erythronate can be enzymatically derived from the PPP intermediate E4P, which suggests that erythronate level can potentially reflect PPP activity. To test this, we manipulated the non-oxidative branch of the PPP, which consists of a series of biochemical transformations catalyzed by transketolase (encoded by genes *TKT, TKTL1*, and *TKTL2* in humans) and transaldolase (encoded by the gene *TALDO1* in humans). It has been reported that the erythronate level in the urine and plasma from transaldolase-deficient patients ^30^ is significantly higher than those in unaffected individuals ^18^. We therefore performed shRNA-mediated silencing of the *TALDO1* gene and found that a 7-fold reduction of its mRNA expression led to an approximately 75% higher erythronate level (**Fig. 4A-B**). The levels of several other PPP intermediates including E4P and upper glycolytic metabolites were also significantly elevated (**Table S1**) in the *TALDO1* knockdown cells with a slight decrease in the transketolase expression (**Fig. 4C**).

**Figure 4.**
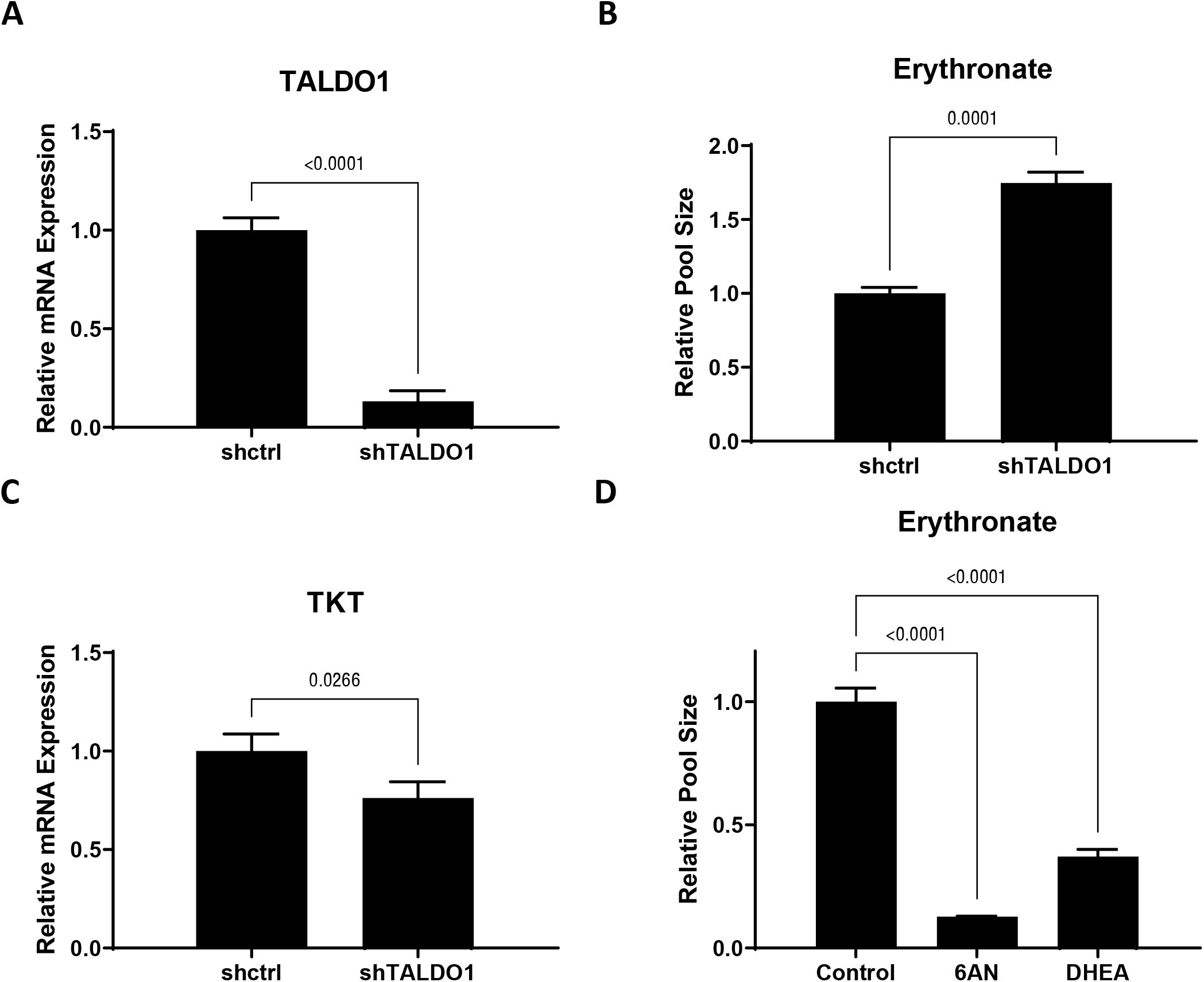
Erythronate level is regulated by enzymes in the pentose phosphate pathway. **A**-**B**) Silencing of *TALDO1* increased the intracellular erythronate level. Erythronate level in *TALDO1* knock-down A549 cells was increased by approximately 70% as compared to the control. **C**) *TALDO1* knock-down led to slightly reduced *TKT* expression. The mRNA level for transaldolase (*TALDO1*) and transketolase (*TKT*) genes in *TALDO1* knock-down A549 cells were determined by real-time quantitative PCR. **D**) Effects of inhibition of the oxPPP on erythronate level in A549 cells. Cells are treated with 6-aminonictinamide (6AN) or *trans*-dehydroepiandrosterone (DHEA), which are known to inhibit enzymes in the oxPPP. The results showed that erythronate level is significantly lowered by the inhibition of oxPPP. Error bars, s.d. (n=3). Statistical test: two-tailed paired t-test (A-C) and one-way ANOVA with Dunnett post-hoc.

To assess whether the oxPPP activity can impact erythronate level, A549 cells were treated with 6-aminonicotinamide (6AN), a drug that strongly inhibits 6-phosphogluconate dehydrogenase (6PGD) and glucose-6-phosphate dehydrogenase (G6PD) ^31^, which are the rate-limiting enzymes of the oxPPP. Culturing A549 cells in DMEM supplemented with 200 µM of 6AN for 1 day resulted in roughly a 10-fold decrease in the erythronate level (**Fig. 4D**). Another G6PD inhibitor, *trans*-dehydroepiandrosterone (DHEA) ^32,33^, had a similar effect on erythronate level in A549 cells (**Fig. 4D**). Presumably, the reduction of erythronate level can be attributed to lower flux through the oxPPP and a corresponding decreased availability of E4P.

### Erythronate level increased in tumor tissues

Having shown the effect of PPP flux on intracellular erythronate levels, we asked whether cancer cells exhibit higher erythronate levels than normal cells, given the observation of higher PPP activity, in various cancer cells ^34–36^. We extracted metabolites from both lung tumors and adjacent normal tissues from 17 lung cancer patients and used GC-MS to assess cancer-induced changes in metabolite levels *in vivo*. Although the level of intracellular erythronate varied among patients, tumor tissue samples from the majority (12 out of 17) showed significantly higher erythronate levels as compared to those from adjacent normal tissues (**Fig. 5**). Only one patient exhibited the opposite trend, where significantly higher erythronate level was detected in healthy tissues.

**Figure 5.**
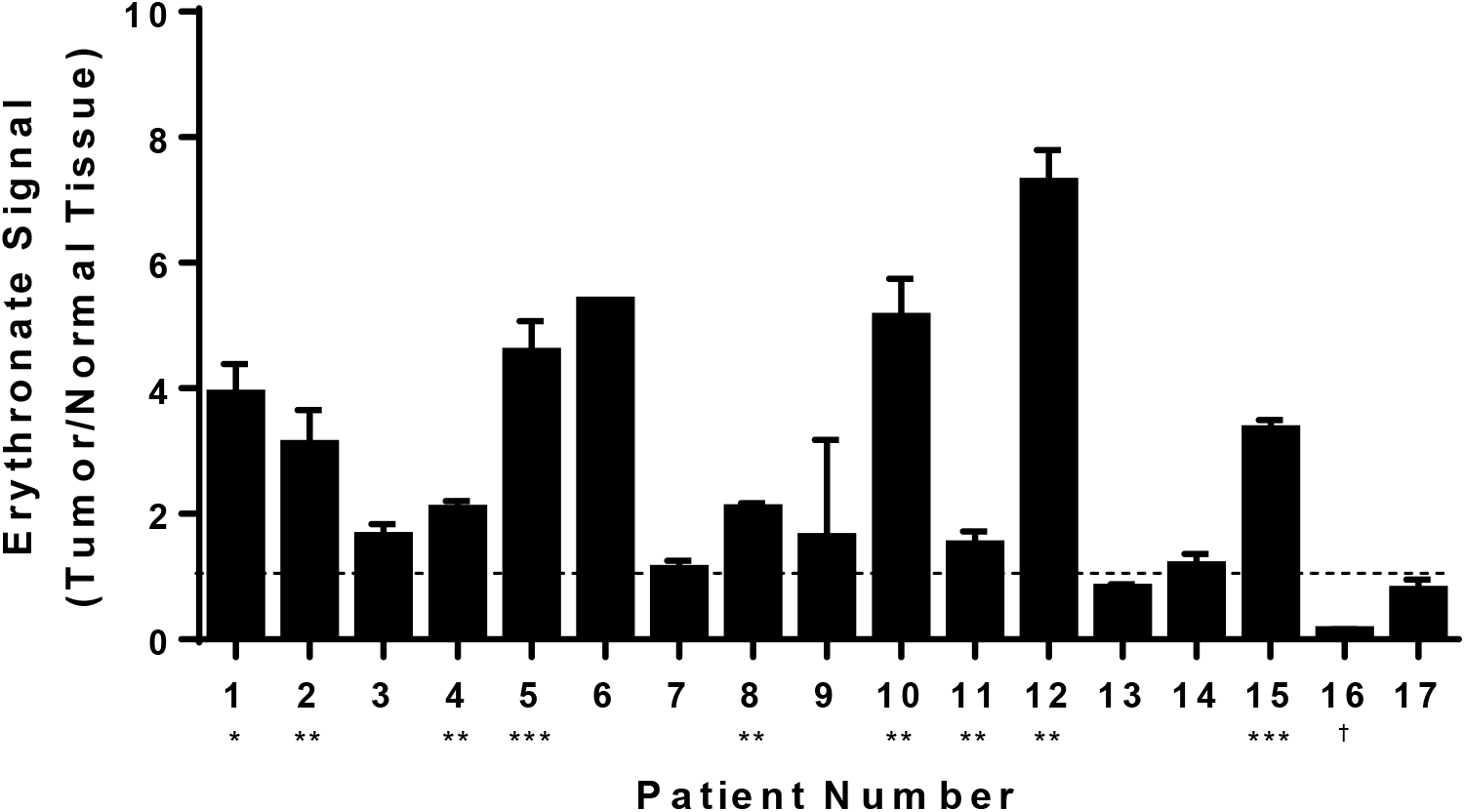
Levels of erythronate in tumor and normal tissues from lung cancer patients. Metabolites were extracted from tissue samples taken from 17 lung cancer patients. GC-MS analysis revealed that, despite various basal levels in healthy tissues from these patients, erythronate (quantified at m/z 292) was significantly increased in tumor tissues as compared to the surrounding normal tissues in 12 of 17 patients. Stars indicate the significance between measurements between tumor and healthy tissues for the same patient. *p-value < 0.05; **p-value < 0.01, ***p-value < 0.001 (Welch’s t-test). † indicates p-value < 0.05 for a decrease in erythronate in tumor tissue relative to normal tissue. Error bars, s.d. (n=3).

## Discussion

Despite the fact that erythronate has been identified to be common to species as varied as fungi to humans ^37,38^, its biosynthetic pathway and biological functions have not been adequately characterized. It has been proposed that erythronate might be formed as a byproduct of the degradation of glycated protein ^39^. However, the time scale of the degradation (several days) does not match the observation that erythronate was formed within minutes in the enzymatic assay using A549 cell lysate (**Fig. 2**). Alternatively, it was speculated that erythronate might be a product of the degradation of ascorbic acid ^38^, but this route is inconsistent with our observations, given that the culture media used in this investigation contained no ascorbic acid. Furthermore, supplementation of 40 mg L^-1^ ascorbic acid in the DMEM medium had no significant effect on the intracellular erythronate level (**Fig. S7**), suggesting that ascorbic acid cannot be the major source of erythronate, if it ever has any role. The finding that erythronate is also produced by the fungi *Phanerochaete chrysosporium* ^40^, *Saccharomyces cerevisiae* and *Yarrowia lipolytica* grown in defined media (data not shown) led us to speculate that erythronate, although barely annotated, is a common metabolite in many (if not all) eukaryotes. Considering that the secondary metabolic pathways of mammals and fungi tend to be very distinct, erythronate is more likely to be derived from central carbon metabolism. More recently, the biosynthesis of erythronate has been elucidated in mammals and yeast. The oxidation and dephosphorylation steps were identified to be catalyzed by GAPDH and PGP ^15,16^. However, this does not exclude the possibility of alternative pathways generating erythronate from E4P.

By performing stable isotopic tracer experiments, we determined that the metabolite erythronate is derived from the third to sixth carbons (C3-6) of glucose. These are the same atoms as those of E4P, implying there might be a route linking E4P and erythronate directly. Two such possibilities, that only differ by the order in which the oxidoreductase and the phosphatase enzymes act, were proposed ^18^. We have shown that A549 cell lysate contains both enzymes required to convert E4P into erythronate, possibly an acid phosphatase and the aldehyde dehydrogenase encoded by *ALDH1A1*. The enzyme responsible for the conversion of E4P to erythronate-4-P, which participates in vitamin B^6^ biosynthesis, exists widely in bacteria but has not been identified in any eukaryotes ^41^. Therefore, it is more likely that a phosphatase first acts on E4P and generates erythrose, which is in turn oxidized by aldehyde dehydrogenase in the presence of NAD^+^. We also found that GAPDH showed very mild activity in converting erythrose into erythronate (**Fig. S5**). This result implies that besides ALDH1A1 and GAPDH, there are likely more promiscuous oxidoreductases that can catalyze the NAD^+^-dependent oxidation of erythrose. Such possibilities include other isoforms of the aldehyde dehydrogenase; however, to identify each of them is beyond the scope of this study.

The identification of ALDH1A1 as the enzyme responsible for erythrose oxidation in cancer cells is interesting, since some cancer cells are known to overexpress various isoforms of aldehyde dehydrogenase, and it was reported that knock-down of *ALDH1A1* and/or another isoform (encoded by *ALDH1A3*) in A549 cells significantly impaired cell growth and motility ^42^. However, it is still unclear by which mechanism high aldehyde dehydrogenase activity promotes cancer growth or if elevated intracellular erythronate has any oncogenic effects. Endogenously-generated aldehydes are cytotoxic at high concentrations ^43^, and we questioned whether elevated erythronate may reflect an enhanced capability of *ALDH1A1*-overexpressing cells to alleviate the toxicity of erythrose, thereby rendering a growth advantage for cancer cells. However, when we cultured empty vector- and *ALDH1A1*-overexpressing MCF-10A cells in 1 mM erythrose, a concentration presumably above physiological conditions, we saw no difference in growth between the two lines (**Fig. S8**). Additionally, cells cultured in erythronate demonstrated no enhancement in proliferation rate. These experiments shed little light on erythronate’s potential functionality, and given biosynthetic pathway outlined in this work, it is possible that its accumulation may simply reflect elevated PPP and ALDH1A1 activity.

It has been reported that erythronate produced by the fungus *Phanerochaete chrysosporium* cultured under 100% oxygen was significantly higher than that under normal air, and the authors speculated that the pathway responsible for erythronate production is sensitive to oxygen stress ^40^. This is consistent with our finding that erythronate is derived from the PPP, given that the oxPPP can be up-regulated in response to oxidative stress or reactive oxygen species ^44^. Indeed, the PPP plays an important role in all proliferating cells on several levels: not only it is the only source of pentoses for the *de novo* biosynthesis of nucleic acids ^45^, but the NADPH generated by the PPP is recognized as a major source of reducing equivalents to fuel anabolic processes (such as *de novo* lipogenesis ^46^), as well as those needed to combat oxidative stress ^47,48^. Evidence suggests that G6PD, the rate-limiting enzyme of the oxPPP, is inhibited by p53, the most frequently mutated tumor suppressor in human tumors ^43^, and that p73, which is frequently overexpressed in human cancers, promotes cell proliferation by increasing G6PD expression ^49,50^. Clinical data of erythronate levels in urine from infants and children showed a clear decrease with age, implying there is a positive correlation of this metabolite with the total proliferation of cells in the body ^37^. This is consistent with our finding that significantly higher levels of erythronate were detected in lung tumor tissues as compared to the normal tissues in patients (**Fig. 5**). Therefore, we propose that erythronate can serve as a target in the investigation of cancer metabolism and diseases related to dysfunction in the PPP, such as transaldolase deficiency, given that erythronate is generally more stable than the other phosphorylated metabolites in the PPP such as E4P, as well as it can be easily detected using GC-MS or LC-MS.

To summarize, through the use of non-targeted and stable-isotope assisted metabolomics we have provided the evidence for a novel metabolic pathway producing erythronate, which is derived from the PPP intermediate E4P through two steps. First, E4P is dephosphorylated by a yet unknown phosphatase, yielding erythrose, which is then oxidized to erythronate by a NAD^+^-dependent aldehyde dehydrogenase (encoded by *ALDH1A1*). Also, the close link between erythronate and PPP makes this metabolite a good target for the investigation of PPP and ALDH1A1 activity, which are both relevant for several diseases including cancer. We also found that erythronate is significantly accumulated in tumor tissues as compared to normal lesions in lung cancer patients, potentially implicating this metabolite as a biomarker for some types of cancer.

### Experimental Procedures

#### Cell culture

All human cell lines were cultured in cell incubators maintained at 37°C and containing 5% CO^2^ as described previously. A549, MCF-10A, HEK-293T, RWPE-1, and RWPE-2 cell lines were obtained from ATCC. Dulbecco’s modified Eagle medium (DMEM; Mediatech) contained 25 mM of D-glucose, 4 mM of L-glutamine, 10% fetal bovine serum (FBS; Sigma) and 100 U mL^-1^ penicillin/streptomycin (Mediatech). For stable isotopic labeling experiments, cells were cultured for 24 h (initially 4×10^5^ cells per well in six-well plates) in DMEM supplemented with a total 25 mM 1:1 (molar ratio) mixture of unlabeled and uniformly-labeled ^13^C D-glucose (Cambridge Isotope Laboratories), 4 mM of L-glutamine and 10% dialyzed FBS. In erythrose toxicity experiments, cells were initially seeded at a density of 1x10^5^ per well in six-well plates, and medium was changed to DMEM either containing or lacking 1 mM erythrose (Sigma) the following day, and total protein content (as a surrogate for total cell number) was harvested following the media change and approximately every 24 h for 3 days. Total protein was extracted using M-PER protein extraction reagent (Thermo Fisher) and quantified using BCA assay (Thermo Fisher) following the manufacturer’s protocol.

#### Enzymatic assay

Whole cell lysate was prepared either using M-PER mammalian protein extraction reagent (Thermo Fisher) or by douncing in hypotonic buffer as previously described with modifications (Mackall 1979). Briefly, A549 cells were grown in DMEM until confluent, and the medium was aspirated and followed by washing with phosphate buffered saline (PBS). Cells were scraped and resuspended after adding 1 ml of ice-cold lysis buffer (10 mM HEPES, 1.5 mM MgCl^2^, 10 mM KCl, 0.5 mM dithiothreitol) and lysed by douncing the suspension 30 times on ice. After centrifugation at >14,000 g and 4°C for 5 min, the supernatant was directly used in the enzymatic assays, which were performed in 96-well micro-titer plates at 37°C unless otherwise specified. The assay mixture contained 5 mM of sodium pyrophosphate and 0.8 mM of NAD^+^ or NADP^+^. In the erythrose oxidation assay, 0.4 mM of erythrose was used. The level of NADH or NADPH was monitored by optical absorbance at 340 nm.

#### Purification and molecular identification of erythrose dehydrogenase

A549 cells were grown to confluency in a 10 cm dish and extracted by scraping into a lysis buffer containing 25 mM HEPES, pH 7.1 and 1x protease inhibitor cocktail (Roche). After three freeze/thaw cycles, the cell extract was incubated for 30 min on ice in the presence of DNAse I (200 U per ml of extract; Roche Applied Science) and 10 mM MgSO^4^. The crude cell extract was centrifuged at 16,000 g and 4°C for 10 min. The supernatant was then loaded at a flow rate of 0.5 ml/min onto an anion exchange column (HiTrap Q HP, 1 ml resin, GE Healthcare) connected to an Äkta purifier system and equilibrated with buffer A (25 mM HEPES, pH 7.0, 1 mM DTT). After a 10-min wash with buffer A, bound proteins were eluted with a 20-min salt gradient (0-2M NaCl in buffer A) at 1 ml/min and 1 ml fractions were collected. Erythrose dehydrogenase activity was measured in the elution fractions by a spectrophotometric assay monitoring NADH production in the presence of erythrose. The active fractions were pooled and desalted on HiTrap desalting columns (GE Healthcare) equilibrated with buffer B (20 mM Tris pH 7.5, 25 mM NaCl). The desalted fractions were used after trypsin digestion for protein sequence identification through LC-MS-MS.

#### Proteomics analysis using mass spectrometry

##### In-gel digest

The purified proteins were separated on an SDS-gel, visualized using a Coomassie Brilliant Blue stain, and the band of interest was cut out of the gel. Proteins were converted to peptides by in-gel digestion with trypsin (Promega) ^51^. The recovered peptides were separated on an in-house packed 15 cm reverse-phase column (3 µm beads, Reprosil, Dr. Maisch HPLC GmbH, Ammerbuch-Entringen, Germany) using a 10-50% acetonitrile linear gradient on an easy-nLC system (Proxeon, Dreieich, Germany). The separated peptides were directly sprayed into a Velos-OrbiTrap mass spectrometer (Thermo Scientific, Dreieich, Germany). The recorded spectra were analyzed using the MaxQuant software package (version 1.2.2.5) ^52^ by matching the data to the IPI human database (version 3.84) with a false discovery rate (FDR) of 1%.

##### In solution digest

Proteins were converted to peptides by a two-step digestion protocol using endopeptidase lys-C (Wako) and trypsin (Promega). The proteins were denatured in denaturing buffer (6 M urea, 2 M thiourea, 20 mM HEPES, pH 8.0) and digested with endopeptidase lys-C for 3 h. The reaction was then diluted four-fold with 50 mM ammonium bicarbonate buffer and 50 U trypsin was added. The reaction was incubated overnight at room temperature. The peptides then underwent the same chromatographic and mass spectral analysis as those produced by the in-gel digest.

##### Gene silencing

Silencing of the gene *TALDO1* was performed as described previously ^53^. Briefly, lentiviral particles carrying either *TALDO1* shRNA (Santa Cruz Biotechnology) or an empty plasmid that only contained a puromycin-resistance gene were first prepared by transfecting 293T cells using FuGENE^®^ 6 Transfection Reagent (Promega). Then A549 cells were cultured in 6-well plates until 80% confluency (8×10^5^ cells per well) and then underwent lentiviral infection. Transfected A549 cells were selected and maintained using DMEM containing 2 mg L^-1^ and 1 mg L^-1^ puromycin, respectively. Silencing of *ALDH1A1* was performed using siRNA (Dharmacon) according to the manufacturer’s protocol. Real time quantitative polymerase chain reaction (RT-qPCR) was performed to determine the change in the target gene expression with reference to *ACTB* (β-actin) as a housekeeping gene. Primers used in the RT-qPCR are listed in **Table S2**.

##### ALDH1A1 overexpression

*ALDH1A1* cDNA was produced through first extracting RNA from A549 cells using a Qiagen RNeasy Plus kit, generating a cDNA library using a Promega GoScript Reverse Transcription System kit, and targeting the processed *ALDH1A1* cDNA for amplification using conventional PCR. Manufacturer’s instructions were followed where applicable. Restriction site-based cloning was used to incorporate *ALDH1A1* cDNA into the pBABE-puro plasmid, and the resulting plasmid was verified through sequencing. pBABE-puro-ALDH1A1 (or negative control empty pBABE-puro) was cotransfected into HEK-293T cells with pUMVC and pCMV-VSV-G plasmids using Promega FuGENE 6 (following manufacturer’s instructions) to produce retroviral particles. Spent 293T medium was harvested after 36 h, filtered through a 0.45 μm-pore membrane, and added with 8 μg/mL polybrene (Millipore) to MCF-10A cells. Cells were incubated with virus for 24 h, and following washing and 24 h in normal medium, selected in 2 μg/mL puromycin (Gemini Bio Products). Cells were cultured in puromycin-containing media until they were used for experiments. RT-qPCR was used to validate *ALDH1A1* overexpression (using *ACTB* as a reference gene). The pBABE-puro plasmid was a gift from Hartmut Land & Jay Morgenstern & Bob Weinberg (Addgene plasmid # 1764) ^54^, and the pUMVC and pCMV-VSV-G plasmids were gifts from Bob Weinberg (Addgene plasmids # 8449 and # 8454, respectively).

##### Metabolomics analysis using GC-MS

###### Metabolite extraction from cell culture

Intracellular metabolites were extracted from cultured human cells using the method described previously with minor modifications ^13^. A549 cells are described as an example: A549 cells were grown in 6-well plate or 6 cm culture dishes until near-confluency. Medium was removed by aspiration, followed by washing with 2 ml of saline. Cells were quenched with 400 µl of -20°C methanol, 300 µl of ice-cold water containing 2 µg norvaline and 2 µg glutarate was added as internal standards, and cells were collected using a cell scraper. The cell suspension in methanol-water solution was then mixed with 400 µl of -20°C chloroform in 1.5 ml tubes. The tubes were shaken vigorously on a vortex mixer at 4°C for 20 min and samples were centrifuged at 14,000×g for 10 min at 4°C. The aqueous phase (methanol-water solution that contained polar metabolites) was collected in a new tube and dried under airflow at room temperature.

###### Metabolite extraction from tissue samples

To investigate the metabolomics differences between cancerous and normal tissues, lung tumor tissue and their surrounding healthy tissue samples were taken from 19 lung cancer patients. The lung tissue was milled (Retch mill) with three steel balls (5 mm diameter) (Conrad Electronics) for 2 min at 22.5 s^-1^. Samples were then homogenized by milling with ice-cold extraction fluid (methanol/water 40:8.5, 485 µl per 100 mg tissue) and five steel balls (1 mm diameter) for 2 min at 22.5 s^-1^. Ice cold water (200 µl per 100 mg tissue) and -20°C chloroform (400 µl per 100 mg tissue) were added, shaken at 1,400 rpm for 20 min at 4°C (Thermomixer Eppendorf) and centrifuged at 5,000×g for 5 min at 4°C. The upper aqueous phase (10-30 ul) was collected in specific glass vials with micro inserts and evaporated under vacuum at -4°C using a refrigerated CentriVap Concentrator (Labconco).

###### Derivatization of polar metabolites

The dried polar metabolites were dissolved and reacted in 12 µl of 2% methoxamine hydrochloride in pyridine (MOX reagent, Thermo) and incubated at 37°C for 1.5 h. The reaction continued after addition of 16 µl of N-Methyl-N-(trimethylsilyl)trifluoroacetamide (MSTFA) + 1% trimethylchlorosilane (TMCS) and incubated at 37°C for 1 h. After derivatization, the samples were centrifuged at 14,000×g for 10 min to remove any precipitation. Alternatively, derivatization was performed using a Gerstel MultiPurpose Sampler (MPS). Dried polar metabolites were dissolved in 15 μl of 2% of MOX reagent at 45°C under shaking. After 30 min, an equal volume of MSTFA was added and held for 30 min at 40°C under continuous shaking. After derivatization, 1 µl of supernatant was injected to GC.

###### GC-MS analysis

GC-MS analysis was performed using an Agilent 6890 or 7890A GC equipped with a 30 m DB-35MS capillary column connected to an Agilent 5975 series MS operating under electron impact ionization at 70 eV. The MS source and quadrupole were held at 230 °C and 150 °C, respectively, and the detector was operated in scan mode. Helium was used as carrier gas at a flow rate of 1 ml min^-1^. The GC oven was operated using the following thermal profile: Hold at 80 °C for 6 min, then increase by 6 °C min^-1^ to 300 °C and hold for 10 min, then increase by 10°C min^-1^ to 325 °C and hold for 4 min. An aliquot of 1 µl C10-40 alkane mixture (Sigma-Aldrich) was run using the same GC program to calculate the retention index for metabolite identification purposes. Identification and quantification of metabolites were performed using the software MetaboliteDetector ^55^. Norvaline-or glutarate-normalized metabolite levels were further normalized to total cell number, protein content, or glutamate ion count (the most abundant target metabolite in GC-MS chromatograms, which exhibited minimal variation when normalized to norvaline/glutarate and cell count/protein content across tested conditions). Isotopically-labeled metabolites were detected and their mass isotopomer distributions were calculated using the software NTFD ^14^.

## Author Contributions

KH and GS conceived the project. JZ, MAK, TC and KH designed the experiments. JZ, MAK, JG, TC, TK, AP, CL, CMM performed experiments. TL provided clinical samples. JZ, MAK, WD, TC, TK, AP, CL, GD and KH analyzed the data. JZ, MAK, WD, TC, KH and GS wrote the manuscript.

## Acknowledgements

Research on cancer metabolism in Stephanopoulos Lab is funded by NIH grants 1R01DK075850-01 and 1R01CA160458-01A1 (USA). J.Z. is supported by a fellowship from LCSB (Luxembourg). This study was supported by the Fonds National de la Recherche, Luxembourg (ATTRACT A10/03). The authors thank Andre Wegner (LCSB) for technical advice in the usage of software MetaboliteDetector and NTFD and fruitful discussion.

## References

1. Hanahan, D. & Weinberg, R. A. Hallmarks of cancer: The next generation. Cell 144, 646–674 (2011).

2. Vander Heiden, M. G. et al. Understanding the Warburg Effect : Cell Proliferation. Science 324, 1029–1034 (2009).

3. Tsouko, E. et al. Regulation of the pentose phosphate pathway by an androgen receptor-mTOR-mediated mechanism and its role in prostate cancer cell growth. Oncogenesis 3, 1–10 (2014).

4. DeBerardinis, R. J. & Chandel, N. S. Fundamentals of cancer metabolism. Sci. Adv. 2, 1–18 (2016).

5. Pavlova, N. N. & Thompson, C. B. The Emerging Hallmarks of Cancer Metabolism. Cell Metab. 23, 27–47 (2016).

6. Pavlova, N. N., Zhu, J. & Thompson, C. B. The hallmarks of cancer metabolism: Still emerging. Cell Metab. 34, 355–377 (2022).

7. Locasale, J. W. et al. Phosphoglycerate dehydrogenase diverts glycolytic flux and contributes to oncogenesis. Nat. Genet. 43, 869–74 (2011).

8. Chaneton, B. et al. Serine is a natural ligand and allosteric activator of pyruvate kinase M2. Nature 491, 458–462 (2012).

9. Ye, J. et al. Pyruvate kinase M2 promotes de novo serine synthesis to sustain mTORC1 activity and cell proliferation. Proc. Natl. Acad. Sci. U. S. A. 109, 6904–6909 (2012).

10. Ward, P. S. et al. The Common Feature of Leukemia-Associated IDH1 and IDH2 Mutations Is a Neomorphic Enzyme Activity Converting α-Ketoglutarate to 2-Hydroxyglutarate. Cancer Cell 17, 225–234 (2010).

11. Hiller, K. & Metallo, C. M. Profiling metabolic networks to study cancer metabolism. Curr. Opin. Biotechnol. 24, 60–68 (2013).

12. Dong, W., Rawat, E. S., Stephanopoulos, G. & Abu-Remaileh, M. Isotope tracing in health and disease. Curr. Opin. Biotechnol. 76, 102739 (2022).

13. Hiller, K., Metallo, C. M., Kelleher, J. K. & Stephanopoulos, G. Nontargeted elucidation of metabolic pathways using stable-isotope tracers and mass spectrometry. Anal. Chem. 82, 6621–6628 (2010).

14. Hiller, K. et al. NTFD--a stand-alone application for the non-targeted detection of stable isotope-labeled compounds in GC/MS data. Bioinformatics 29, 1226–1228 (2013).

15. Ishii, Y., Hashimoto, T., Minakami, S. & Yoshikawa, H. The formation of erythronic acid 4-phosphate from erythrose 4-phosphate by glyceraldehyde-3-phosphate dehydrogenase. J. Biochem. 56, 111–112 (1964).

16. Collard, F. et al. A conserved phosphatase destroys toxic glycolytic side products in mammals and yeast. Nat. Chem. Biol. 12, 601–607 (2016).

17. Ishii, Y. Erythronic acid 4-phosphate, a new intermediate of inosine metabolism in human red cell hemolysate. J. Biochem. 55, 371–377 (1964).

18. Engelke, U. F. H. et al. Mitochondrial involvement and erythronic acid as a novel biomarker in transaldolase deficiency. Biochim. Biophys. Acta. 1802, 1028–1035 (2010).

19. Perl, A., Hanczko, R., Telarico, T., Oaks, Z. & Landas, S. Oxidative stress, inflammation and carcinogenesis are controlled through the pentose phosphate pathway by transaldolase. Trends Mol. Med. 17, 395–403 (2011).

20. Jonas, S. & Hollfelder, F. Mapping catalytic promiscuity in the alkaline phosphatase superfamily. Pure Appl. Chem. 81, 731–742 (2009).

21. López-Canut, V., Roca, M., Bertrán, J., Moliner, V. & Tuñón, I. Promiscuity in alkaline phosphatase superfamily. Unraveling evolution through molecular simulations. J. Am. Chem. Soc. 133, 12050–12062 (2011).

22. Anand, A. & Srivastava, P. K. A molecular description of acid phosphatase. Appl. Biochem. Biotechnol. 167, 2174–2197 (2012).

23. Marcato, P. et al. Aldehyde dehydrogenase activity of breast cancer stem cells is primarily due to isoform ALDH1A3 and its expression is predictive of metastasis. Stem Cells 29, 32–45 (2011).

24. Ginestier, C. et al. ALDH1 Is a Marker of Normal and Malignant Human Mammary Stem Cells and a Predictor of Poor Clinical Outcome. Cell Stem Cell 1, 555–567 (2007).

25. Landen, C. N. et al. Targeting aldehyde dehydrogenase cancer stem cells in ovarian cancer. Mol. Cancer Ther. 9, 3186–3199 (2010).

26. Bello, D., Webber, M. M., Kleinman, H. K., Wartinger, D. D. & Rhim, J. S. Androgen responsive adult human prostatic epithelial cell lines immortalized by human papillomavirus 18. Carcinogenesis 18, 1215–1223 (1997).

27. Li, X. shan, Xu, Q., Fu, X. yang & Luo, W. sheng. ALDH1A1 overexpression is associated with the progression and prognosis in gastric cancer. BMC Cancer 14, 1–8 (2014).

28. Xu, S. L. et al. Aldehyde dehydrogenase 1A1 circumscribes high invasive glioma cells and predicts poor prognosis. Am. J. Cancer Res. 5, 1471–1483 (2015).

29. Sacco, F., Perfetto, L., Castagnoli, L. & Cesareni, G. The human phosphatase interactome: An intricate family portrait. FEBS Letters 586, 2732–2739 (2012).

30. Verhoeven, N. M. et al. Transaldolase deficiency: Liver cirrhosis associated with a new inborn error in the pentose phosphate pathway. Am. J. Hum. Genet. 68, 1086–1092 (2001).

31. Dong, W. et al. Oncogenic metabolic rewiring independent of proliferative control in human mammary epithelial cells. bioRxiv 2022, https://doi.org/10.1101/2022.04.08.486845.

32. Tian, W. N. et al. Importance of glucose-6-phosphate dehydrogenase activity for cell growth. J. Biol. Chem. 273, 10609–10617 (1998).

33. Shantz, L. M., Talalay, P. & Gordon, G. B. Mechanism of inhibition of growth of 3T3-L1 fibroblasts and their differentiation to adipocytes by dehydroepiandrosterone and related steroids: Role of glucose-6-phosphate dehydrogenase. Proc. Natl. Acad. Sci. U. S. A. 86, 3852–3856 (1989).

34. Langbein, S. et al. Metastasis is promoted by a bioenergetic switch: New targets for progressive renal cell cancer. Int. J. Cancer 122, 2422–2428 (2008).

35. Chen, Z. et al. Overexpression of G6PD is associated with poor clinical outcome in gastric cancer. Tumor Biol. 33, 95–101 (2012).

36. Van Driel, B. E. M. et al. Prognostic estimation of survival of colorectal cancer patients with the quantitative histochemical assay of G6PDH activity and the multiparameter classification program CLASSIF1. Cytometry 38, 176–183 (1999).

37. Guneral, F. & Bachmann, C. Age-related reference values for urinary organic acids in a healthy Turkish pediatric population. Clin. Chem. 40, 862–866 (1994).

38. Harding, J. J., Hassett, P., Rixon, K. C., Bron, A. J. & Harvey, D. J. Sugars including erythronic and threonic acids in human aqueous humour. Curr. Eye Res. 19, 131–136 (1999).

39. Ahmed, M. U., Thorpe, S. R. & Baynes, J. W. Identification of N(ε)-carboxymethyllysine as a degradation product of fructoselysine in glycated protein. J. Biol. Chem. 261, 4889–4894 (1986).

40. Miura, D., Tanaka, H. & Wariishi, H. Metabolomic differential display analysis of the white-rot basidiomycete Phanerochaete chrysosporium grown under air and 100% oxygen. FEMS Microbiol. Lett. 234, 111–116 (2004).

41. Lee, J. Y. et al. Overexpression, crystallization and preliminary X-ray crystallographic analysis of the variable lymphocyte receptor 2913 ectodomain fused with internalin B. Acta Crystallogr. Sect. F Struct. Biol. Cryst. Commun. 69, 39–41 (2013).

42. Moreb, J. S. et al. ALDH isozymes downregulation affects cell growth, cell motility and gene expression in lung cancer cells. Mol. Cancer 7, 1–19 (2008).

43. O’Brien, P., Siraki, A. & Shangari, N. Aldehyde sources, metabolism, molecular toxicity mechanisms, and possible effects on human health. Crit. Rev. Toxicol. 35, 609–662 (2005).

44. Salvemini, F. et al. Enhanced glutathione levels and oxidoresistance mediated by increased glucose-6-phosphate dehydrogenase expression. J. Biol. Chem. 274, 2750–2757 (1999).

45. Ahn, W. S. et al. Glyceraldehyde 3-phosphate dehydrogenase modulates nonoxidative pentose phosphate pathway to provide anabolic precursors in hypoxic tumor cells. AIChE J. 64, 4289–4296 (2018).

46. Park, J. et al. Overexpression of Glucose-6-Phosphate Dehydrogenase Is Associated with Lipid Dysregulation and Insulin Resistance in Obesity. Mol. Cell. Biol. 25, 5146–5157 (2005).

47. Cosentino, C., Grieco, D. & Costanzo, V. ATM activates the pentose phosphate pathway promoting anti-oxidant defence and DNA repair. EMBO J. 30, 546–555 (2011).

48. Moon, S. J., Dong, W., Stephanopoulos, G. N. & Sikes, H. D. Oxidative pentose phosphate pathway and glucose anaplerosis support maintenance of mitochondrial NADPH pool under mitochondrial oxidative stress. Bioeng. Transl. Med. 5, 1–18 (2020).

49. Amelio, I. et al. P73 regulates serine biosynthesis in cancer. Oncogene 33, 5039–5046 (2014).

50. Du, W. et al. TAp73 enhances the pentose phosphate pathway and supports cell proliferation. Nat. Cell Biol. 15, 991–1000 (2013).

51. Shevchenko, A., Tomas, H., Havliš, J., Olsen, J. V. & Mann, M. In-gel digestion for mass spectrometric characterization of proteins and proteomes. Nat. Protoc. 1, 2856–2860 (2007).

52. Cox, J. & Mann, M. MaxQuant enables high peptide identification rates, individualized p.p.b.-range mass accuracies and proteome-wide protein quantification. Nat. Biotechnol. 26, 1367–1372 (2008).

53. Metallo, C. M. et al. Reductive glutamine metabolism by IDH1 mediates lipogenesis under hypoxia. Nature 481, 380–384 (2012).

54. Morgenstern, J. P. & Land, H. Advanced mammalian gene transfer: High titre retroviral vectors with multiple drug selection markers and a complementary helper-free packaging cell line. Nucleic Acids Res. 18, 3587–3596 (1990).

55. Hiller, K. et al. Metabolite detector: Comprehensive analysis tool for targeted and nontargeted GC/MS based metabolome analysis. Anal. Chem. 81, 3429–3439 (2009).

56. Dong, W., Moon, S. J., Kelleher, J. K. & Stephanopoulos, G. Dissecting Mammalian Cell Metabolism through 13C-And 2H-Isotope Tracing: Interpretations at the Molecular and Systems Levels. Ind. Eng. Chem. Res. 59, 2593–2610 (2020).

