## Supplemental Materials for "Non-targeted metabolomics identifies erythronate accumulation in cancer cells"

Cancer metabolism; non-targeted metabolomics; erythronate; pentose phosphate pathway

**Table S1. Change of PPP and glycolytic intermediates caused by knockdown of TALDO1**

| Metabolite | [TALDO1 KD]/[Control] | p-value (student t-test, n=4) |
|---|---|---|
| Fructose 1,6-biP | 1.93 | 0.007 |
| Glucose 6-P | 2.07 | 0.001 |
| Ribose 5-P | 1.74 | 0.0006 |
| Ribulose 5-P | 2.06 | 0.0001 |
| Xylulose 5-P | 1.44 | 0.0007 |
| Erythrose 4-P | 3.12 | 0.01 |

**Table S2. Primers used in the RT-qPCR**

| Primer | Sequence 5’ -> 3’ | Location (bp) |
|---|---|---|
| ACTB_Fw | CATGTACGTTGCTATCCAGGC | 393-413 |
| ACTB_Rv | CTCCTTAATGTCACGCACGAT | 642-622 |
| TALDO1_Fw | CTCACCCGTGAAGCGTCAG | 9-27 |
| TALDO1_Rv | GTTGGTGGTAGCATCCTGGG | 135-116 |
| TKT_Fw | TCCACACCATGCGCTACAAG | 158-177 |
| TKT_Rv | CAAGTCGGAGCTGATCTTCCT | 321-301 |
| ALDH1A1_Fw | GCACGCCAGACTTACCTGTC | 14-33 |
| ALDH1A1_Rv | CCTCCTCAGTTGCAGGATTAAAG | 142-120 |

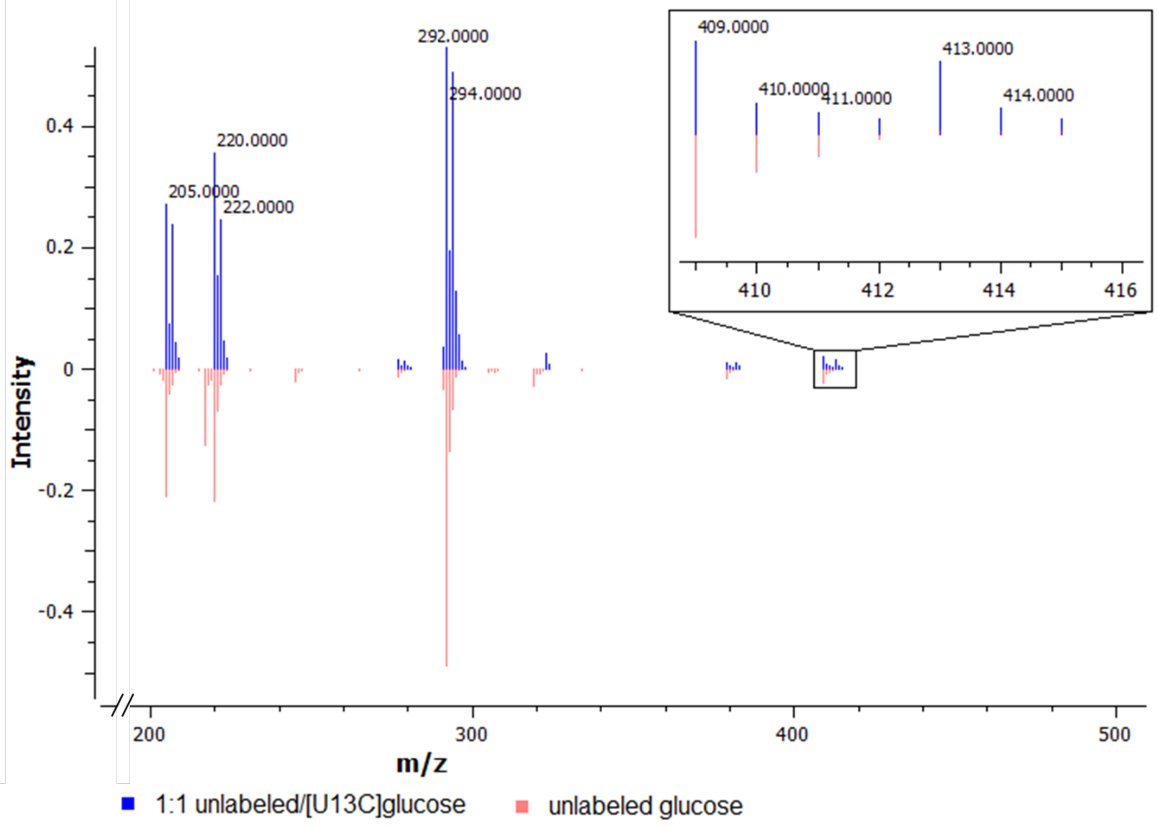


**Figure S1. Labeling of erythronate by [U-$^{13}C_6$]glucose in cancer cells.**

A549 lung carcinoma cells were cultured with media containing either unlabeled glucose (red) or a 1:1 mixture of unlabeled and [U-$^{13}C_6$]glucose (blue). Intracellular polar metabolites were extracted from the cells and analyzed on GC-MS. Note the abundance shift at m/z 413 indicates the M+4 mass isotopomer of erythronate.

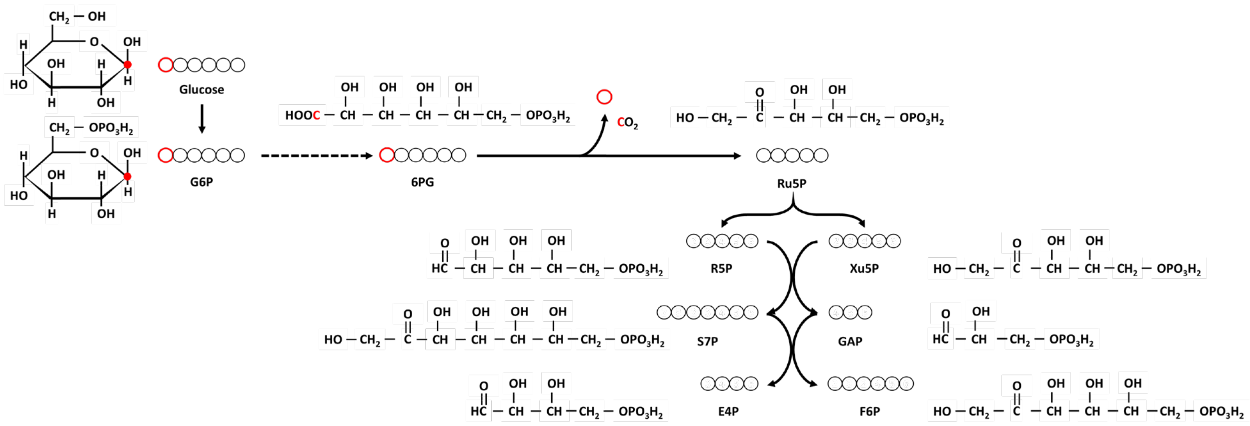


**Fig S2. Chemical structures and reactions along the pentose phosphate pathway (PPP).**

This figure is adapted from Dong et al., 2019.

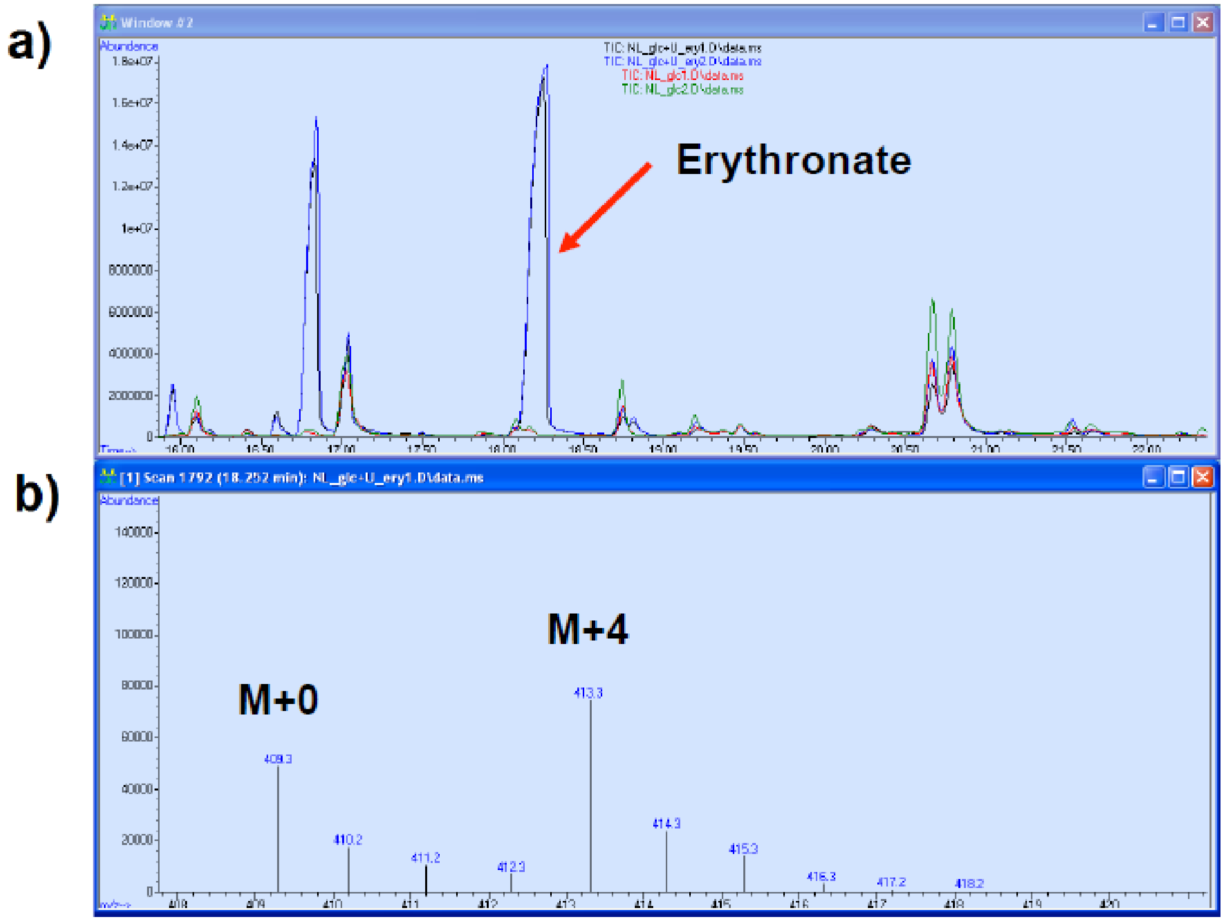


**Figure S3. Erythronate produced in A549 cell culture fed with labeled erythrose.**

a) A549 cells were cultured in DMEM media containing either no erythrose (green and red) or 1 g $L^{-1}$ of [U-$^{13}C_4$]erythrose (black and blue). Large amount of erythronate, observed as the largest peak at around 18 min, was produced by A549 cells, meaning erythrose can cross cell membrane and be oxidized to erythronate in the cell; b) Mass spectrum of ion 409 of TMS-derivatized erythronate shows that erythronate was M+4 labeled when A549 cells were fed with [U-$^{13}C_4$]erythrose.

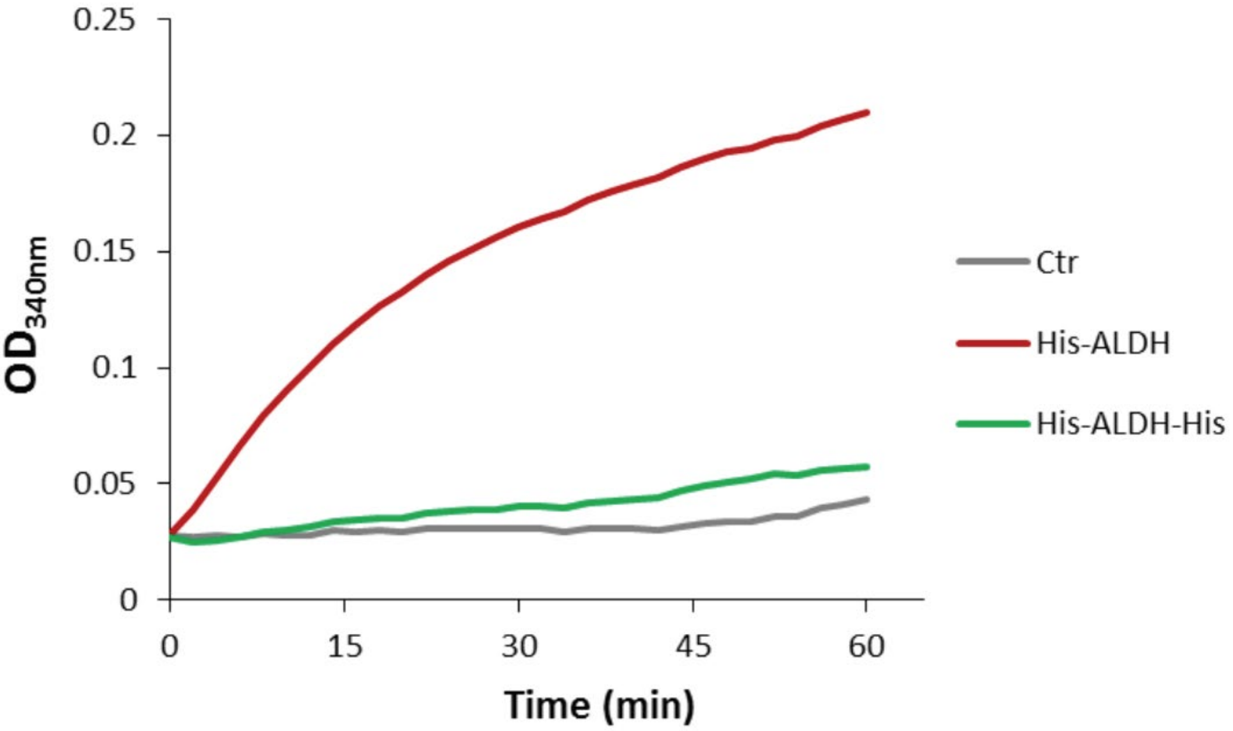


**Figure S4. Enzymatic assay of erythrose oxidation by purified ALDH1A1**.

ALDH1A1 cDNA was reverse transcribed from A549 total mRNA and cloned into a pET28A bacterial expression vector under a T7 promoter. A polyhistidine tag was fused to the N-terminus (red line) or both termini (green) of the protein and used for purification. A blank pET28A vector was used as control. Equal amount of purified proteins (His-Aldh and His-Aldh-His) and the control were used in enzymatic assay for conversion of erythrose to erythronate in the presence of $NAD^+$. The reaction was monitored by absorbance at 340 nm. The results showed that Aldh with polyhistidine tagged on the N-terminus, but not the one tagged on both termini, was able to oxidize erythrose.

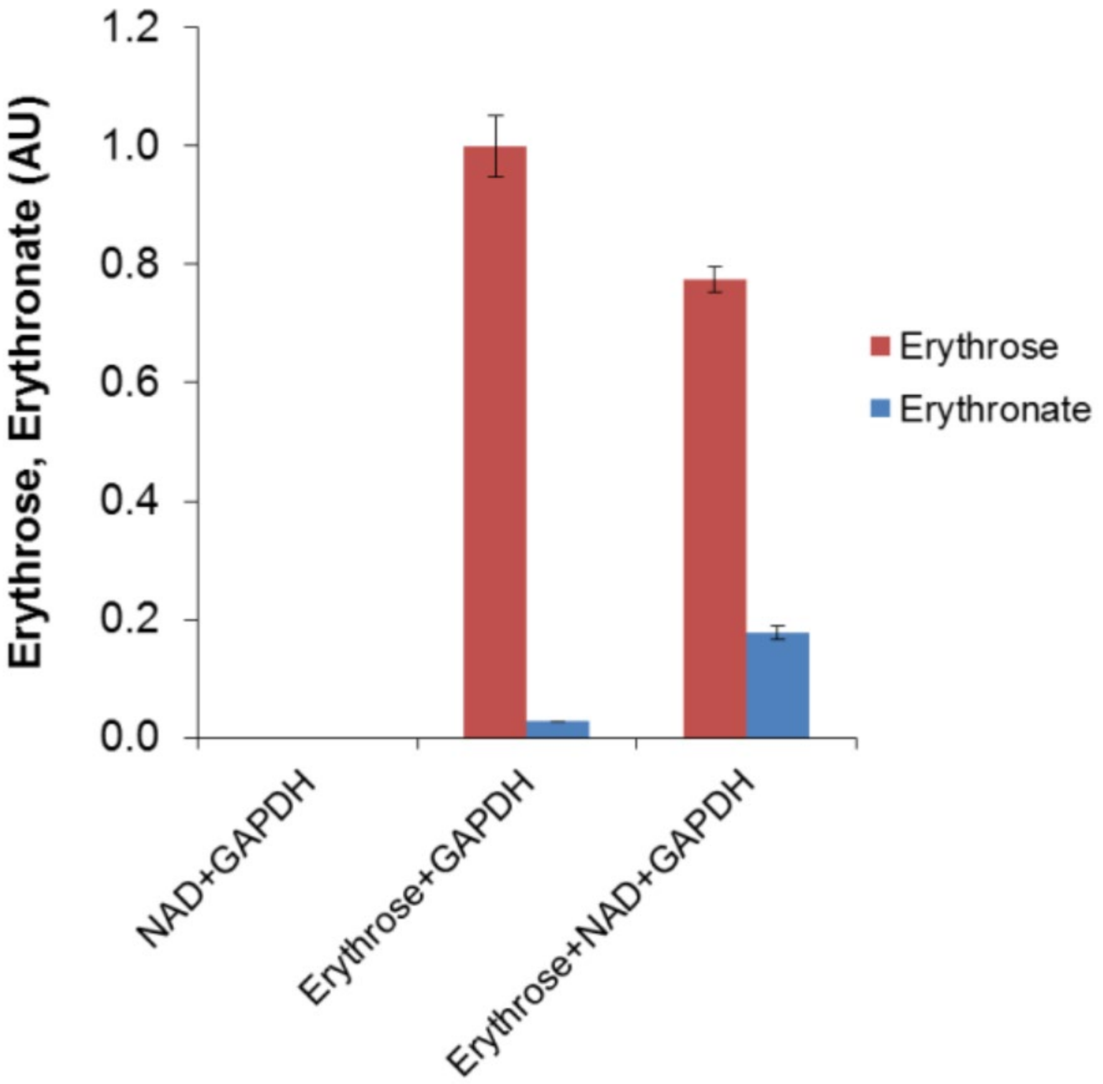


**Figure S5. Conversion of erythrose to erythronate by GAPDH.**

GAPDH was used in the enzymatic assay for the conversion of erythrose to erythronate. The results showed that GAPDH had a minor erythrose-oxidizing activity, and the conversion was enhanced by the addition of 0.8 mM $NAD^{+}$.

A)

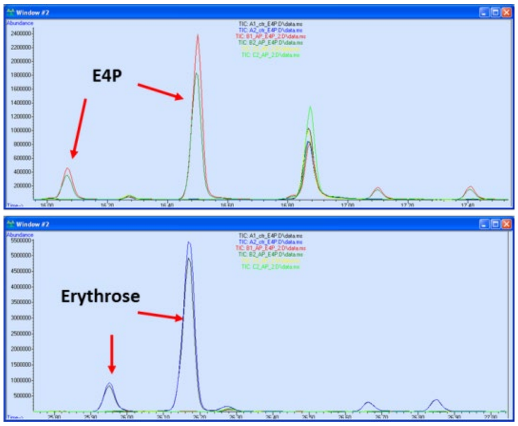


B)

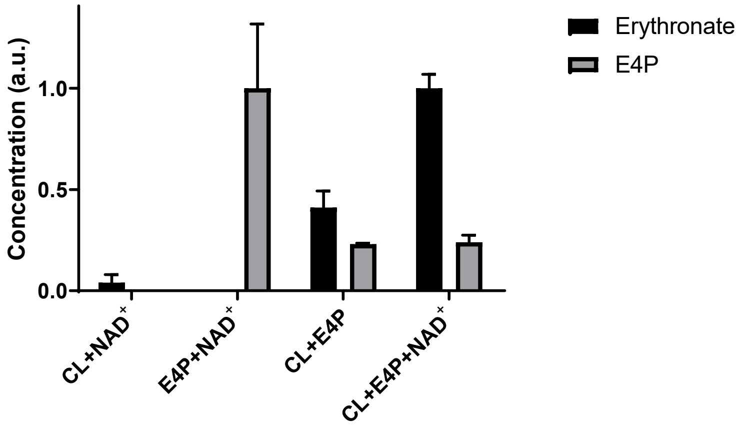


**Figure S6. Conversion of E4P to erythronate by A549 cell lysate. A**) Dephosphorylation of E4P by acid phosphatase (AP). Enzymatic assays were performed in duplicates (represented by lines with different color). **B**) Enzymatic assays were performed using A549 cell lysate (CL) and E4P, in the absence and presence of $NAD^+$. Assay mixtures were analyzed using GC-MS to quantify the relative amounts of E4P and erythronate. The results showed that A549 cell lysate is capable of converting E4P to erythronate and that the conversion is enhanced by the addition of 0.8 mM $NAD^+$. Note that instrumental responses are different for erythronate and E4P, rendering disproportional changes for the target analytes.

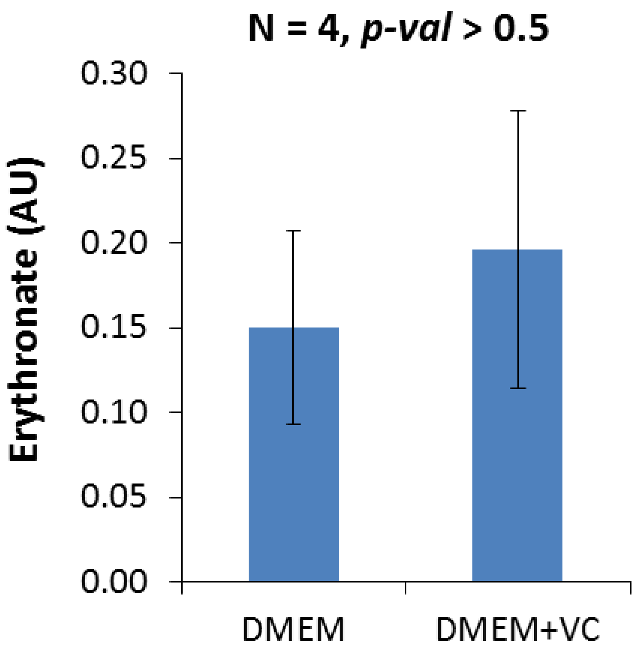


**Figure S7. Effect of ascorbic acid supplementation in DMEM medium on the level of erythronate in A549 cells.**

A549 cells were cultured in normal DMEM medium or supplemented with 40 mg $L^{-1}$ ascorbic acid. Intracellular polar metabolites were extracted and analyzed by GC-MS. Results showed no significant difference between cells grown in DMEM with or without ascorbic acid supplementation.

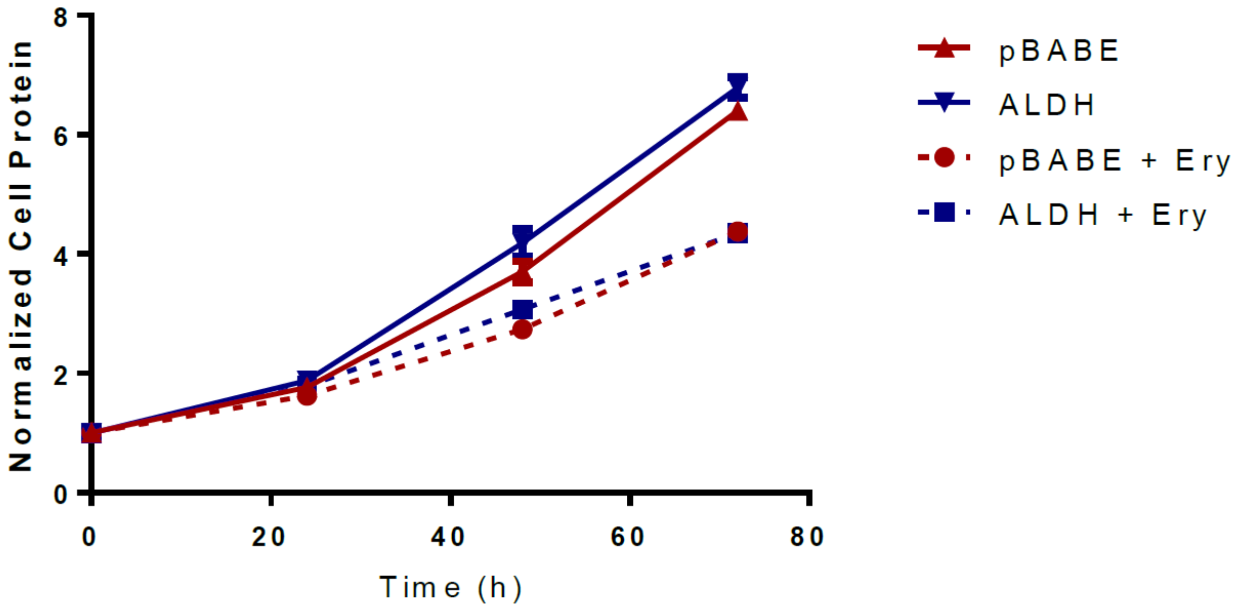


**Figure S8. Growth of ALDH1A1-overexpressing cells in erythrose.**

Empty vector- (pBABE) and ALDH1A1-overexpressing (ALDH) MCF-10A cells were cultured in media with and without 1 mM erythrose (Ery) to monitor growth under erythrose-detoxifying conditions.
